# Edge fires drive the shape and stability of tropical forests

**DOI:** 10.1101/265124

**Authors:** Laurent Hébert-Dufresne, Adam F. A. Pellegrini, Uttam Bhat, Sidney Redner, Stephen W. Pacala, Andrew M. Berdahl

**Author notes:** **Corresponding author**: Andrew Berdahl, Santa Fe Institute, 1399 Hyde Park Rd, Santa Fe, New Mexico, USA, 87501 ☎, ₗ.

## Abstract

In tropical regions, fires propagate readily in grasslands but typically consume only edges of forest patches. Thus forest patches grow due to tree propagation and shrink by fires in surrounding grasslands. The interplay between these competing edge effects is unknown, but critical in determining the shape and stability of individual forest patches, as well the landscape-level spatial distribution and stability of forests. We analyze high-resolution remote-sensing data from protected areas of the Brazilian Cerrado and find that forest shapes obey a robust perimeter-area scaling relation across climatic zones. We explain this scaling by introducing a heterogeneous fire propagation model for tropical forest-grassland ecotones. Deviations from this perimeter-area relation determine the stability of individual forest patches. At a larger scale, our model predicts that the relative rates of tree growth due to propagative expansion and long-distance seed dispersal determine whether collapse of regional-scale tree cover is continuous or discontinuous as fire frequency changes.

## INTRODUCTION

In a variety of ecosystems, a small change in a driving parameter can result in a dramatic shift between disparate regimes. Prominent examples include the eutrophication of lakes (Conley et al., 2009; Smith and Schindler, 2009), the degradation of coral reefs (Mumby et al., 2007), and the collapse of forests (Cochrane et al., 1999; Scheffer et al., 2001). In the particular case of tropical rainforests, it is hypothesized that an abrupt shift between grass- and forest-dominated regimes is due to a fire-mediated bistability (Dantas et al., 2016; Hirota et al., 2011; Staver et al., 2011). This recent work has demonstrated a bistability at the macroscale and highlights the importance of fire as a feedback mechanism that leads to this bistability. However, vegetation-fire feedbacks are inherently local spatial processes (Laurance et al., 2002; Laurance and Yensen, 1991). Incorporating such local processes may help understand how the size and shape of individual forest fragments determine patterns in, and the stability of, the composite system at the regional scale.

In tropical regions, fires that start in grassland typically burn only the edges of forested regions, but do not spread within the forest (Brinck et al., 2017; Cochrane and Laurance, 2002; Hoffmann et al., 2012b; Laurance et al., 2002; Uhl and Kauffman, 1990). This behavior starkly contrasts with wildfires in temperate forests, where the fire spreads readily through the trees via local ignition of neighboring trees (Romme and Despain, 1989; Turner, 2005). This characteristic of fire propagation in the tropics suggests that the ratio of the perimeter of a forested patch (trees adjacent to grassland) to its area (i.e., its protected interior) determines the relative vulnerability of such a patch to fire. On the other hand, the perimeter also determines the potential for the outward expansion of a forested patch due to new tree growth at the perimeter. Thus the stability of an individual forest patch must depend on its *shape* as a result of the interplay between recession due to exposure to fire and expansion due to perimeter-mediated growth.

At a larger scale, in Neotropical systems, burning has led to large-scale forest fragmentation so that areas that were once covered by forest subsequently consist of disjoint forest patches that are interspersed with grassy areas; we term this type of morphology as savanna (Cochrane, 2003; Cochrane et al., 1999; Nepstad et al., 1999; Pueyo et al., 2010). The state of the system is then driven by the collective fate of its constituent forest patches. Hence, edge effects have the potential to drive the dynamics of forest patches at all scales: from the collapse of individual patches to large-scale regime shifts between forest and savanna. While it has been shown that harsher conditions at patch edges are deleterious to trees and increase the trees’ tendency to burn (Laurance et al., 2002; Laurance and Yensen, 1991; Uhl and Kauffman, 1990), the way such edge effects scale up to determinate the shape and stability of forest patches is unknown.

In this work, we investigate the mechanisms that govern the size and shapes of forests in the Brazilian Cerrado, and attempt to understand how perturbations in the shape of a forest can affect its stability. To do so, we first examine high-resolution satellite imagery (∼ 30 × 30m resolution (Hansen et al., 2013)) of savanna-forest ecotones in the Cerrado to quantify the statistical properties of the forested areas. We find a robust scaling relation between the perimeter, *P*, and area, *A*, of forest patches, in which *P* ∼ *A^γ^* with *γ* ≈ 0.69.

To explain these observations, we introduce a mechanistic forest-fire model that is specifically tailored to account for the particular phenomenology of fire propagation in tropical regions that consist of both forested and grassy areas. In our model, fires spread readily through grass but less readily through forested areas. We denote this heterogeneously burning grass and trees process as the BGT model. The spatial version of the BGT model reproduces the same perimeter to area scaling relationship that we observe in the empirical data. We also show that deviations from this area-perimeter relation can be used to infer the stability of individual forest patches and thus suggest what interventions might lead to, or prevent, forest fragmentation and the collapse of forest cover. These local spatial features appear remarkably robust to changes in model parameters.

Finally, we observe interesting macro-scale (>> 30m) behavior in both simulations of our spatial BGT model and numerical solutions of its mean-field version. Specifically, both models exhibit a cusp bifurcation (Kuznetsov, 2013) in total tree cover, in which both discontinuous (first-order) or continuous (second-order) transitions observed between forested and savanna states as a function of model parameters. In the former case, microscopic perturbations to parameters can lead to macroscopic changes in the amount of forest cover.

## MATERIAL AND METHODS

### Data Collection

The Brazilian Cerrado contains a mosaic of grassy areas and forests that are interspersed throughout the landscape (Fig. 1A-C). We downloaded tree cover data for the year 2000 (Hansen et al. (2013), available from: http://earthenginepartners.appspot.com/science-2013-global-forest) and analyzed the subset of this data that corresponds to 86 Category-II protected areas in the Brazilian Cerrado (IUC, 2016), corresponding to a total area of 5.16 million hectares. We focused on the Cerrado because of its mixed forest-savanna ecotone and specifically IUCN- designated protected areas within it to assess fire dynamics in forests and grasslands where there has been no deliberate human intervention. The resolution of the data is 30 × 30m and each cell is characterized by a continuous tree-cover variable (Hansen et al., 2013), from which we assign each cell as either forest when it contains > 50% tree cover or grass when it contains < 50% tree cover. The distribution of tree cover in these cells follows a roughly bimodal distribution, with peaks at very high and very low tree cover, see Appendix Fig. A.2. Consequently, our definitions for forest and grass patches are not sensitive to minor changes in the threshold used. As a check, we performed a sensitivity analysis to our assumption of a 50% threshold, and replicated the analysis using a 75% tree cover threshold.

**Figure 1:**
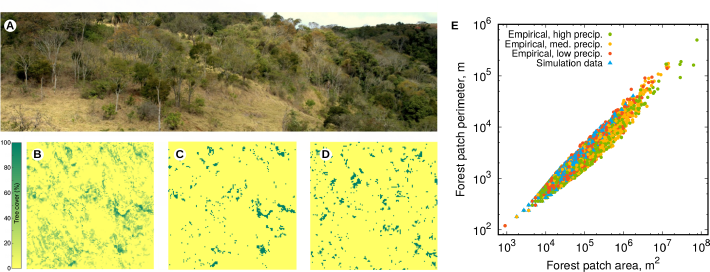
Forest shapes. **(A)** Mixed forest-grassland ecosystem characteristic of the study region, the Brazilian Cerrado. **(B)** Raw forest cover percentage inferred from the high-resolution remote-sensing data, over a 4 km × 4 km area. **(C)** Binary grass-forest state obtained using a 50% threshold on the data of panel (B). **(D)** Snapshot of our spatial model, with same color scheme and spatial scale as (C). **(E)** Perimeter versus area of individual forest patches. The circles denote empirical data (separated in three classes according to precipitation levels) while the triangles show one set of simulation data (see Table 1 for parameter values).

The results are reported below and consistent with those found using the 50% threshold (Table 1).

We then combined forest cells that shared an edge into discrete forest patches, and discarded any patches that were not fully contained within the area of interest since the perimeters of these boundary patches would be artificially short. From this data, we then calculated the area and perimeter length of each forest patch.

**Table 1:** Summary of scaling results for simulations and empirical data.

| | Description | Lattice size | Time step | $p_\lambda$ | $p_\alpha$ | $p_\beta$ | $p_f$ | $p_G$ | $p_T$ | $p_\mu$ | Scaling | Confidence interval | Presented |
| --- | --- | --- | --- | --- | --- | --- | --- | --- | --- | --- | --- | --- | --- |
| Simulations | low $\alpha/\beta$ ratio | 768 | one year | 1.0 | 0.03 | 0.0003 | $6.2 \cdot 10^{-7}$ | 0.9 | 0.1 | 1.0 | 0.68947 | [0.68205 0.69698] | |
| | medium $\alpha/\beta$ ratio | 768 | one year | 1.0 | 0.03 | 0.001 | $7.0 \cdot 10^{-7}$ | 0.9 | 0.1 | 1.0 | 0.71111 | [0.70150 0.72083] | Fig. 1 |
| | alternate lattice | 512 | one year | 1.0 | 0.03 | 0.001 | $1.6 \cdot 10^{-6}$ | 0.9 | 0.1 | 1.0 | 0.70690 | [0.69718 0.71672] | Fig. 2 |
| | alternate fire spread | 512 | one year | 1.0 | 0.03 | 0.0003 | $7.0 \cdot 10^{-7}$ | 0.8 | 0.2 | 1.0 | 0.68639 | [0.67746 0.69541] | |
| | alternate time step | 512 | 1/10 year | 0.1 | 0.004 | $10^{-5}$ | $4.0 \cdot 10^{-7}$ | 0.9 | 0.1 | 1.0 | 0.66622 | [0.66077 0.67170] | |
| | high $\alpha/\beta$ ratio 1 | 256 | 1/10 year | 0.1 | 0.022 | $1.0 \cdot 10^{-6}$ | $3.4 \cdot 10^{-6}$ | 0.9 | 0.1 | 1.0 | 0.66914 | [0.64761 0.69130] | Appendix Fig. C4 |
| | high $\alpha/\beta$ ratio 2 | 256 | 1/10 year | 0.1 | 0.0022 | $2.0 \cdot 10^{-6}$ | $3.9 \cdot 10^{-6}$ | 0.9 | 0.1 | 1.0 | 0.69191 | [0.68228 0.70166] | |
| Empirical | 50% all data | — | — | — | — | — | — | — | — | — | 0.69167 | [0.68234, 0.70111] |  |
|  | 75% all data | — | — | — | — | — | — | — | — | — | 0.68382 | [0.67577, 0.69195] |  |
|  | 50% low precip. | — | — | — | — | — | — | — | — | — | 0.69630 | [0.67682, 0.71623] | Fig. 1 |
|  | 50% med. precip. | — | — | — | — | — | — | — | — | — | 0.70435 | [0.69262, 0.71625] | Fig. 1 |
|  | 50% high precip. | — | — | — | — | — | — | — | — | — | 0.67826 | [0.65163, 0.70585] | Fig. 1 |
|  | 75% low precip. | — | — | — | — | — | — | — | — | — | 0.70748 | [0.69649, 0.71861] |  |
|  | 75% med. precip. | — | — | — | — | — | — | — | — | — | 0.70000 | [0.68601, 0.71424] |  |
|  | 75% high precip. | — | — | — | — | — | — | — | — | — | 0.65833 | [0.63144, 0.68620] |  |

### Spatial BGT Model

We are interested in understanding how the feedback between fire and the regrowth of vegetation determines the distribution of forest shapes and how the geometrical shape of a forest determines its stability. To this end, we formulate a spatial version of our BGT model that incorporates the basic features that forest patches primarily grow outward at their perimeters and burn exclusively at their edges. The four fundamental aspects of our model that parsimoniously account for tropical forest-grassland fire dynamics are: (a) trees spread both locally at the perimeter and spontaneously due to seed dispersal; (b) fires sporadically start in grassland but not in forests; (c) fires percolate readily in grassland; (d) fires percolate only a short distance into forested areas (Cochrane and Laurance, 2002; Hoffmann et al., 2012b; Laurance et al., 2002). This model is similar to, yet more parsimonious than, other cellular automata (CA) forest-fire models designed for tropical forest-savanna systems (Berjak and Hearne, 2002; Favier et al., 2004; Yassemi et al., 2008); however, it contrasts with CA forest fire models from the physics literature (Bak et al., 1990; Chen et al., 1990; Clar et al., 1996; Drossel and Schwabl, 1992; Grassberger, 2002) in that it incorporates heterogeneous burning between grass and trees, and both propagative and dispersed tree growth. These are the key features that allow us to look at forest shapes in a novel way.

We discretize the land area as a two-dimensional lattice (with periodic boundary conditions) where sites interact with their four nearest neighbors. Each site can be occupied by trees, by grass, by ash/dirt, or by burning material— either burning grass or trees. At each time step, the densities of tree, grass, ash, and burning sites, T, G, A, and B evolve according to the following processes:

1. Grass growth: An ash site turns to a grass site with probability *p_λ_.*
2. Tree spreading: A tree site converts a *neighboring* grass or ash site into another tree site with probability *p_α_.*
3. Spontaneous tree growth: Any grass or ash site can turn into a tree site with a small probability *p_β_.* Such a process arises from the long-distance dispersal of tree seeds.
4. Fire ignition and propagation: A grass site ignites with probability *p_f_*. Once a fire starts, it spreads as follows: (a) grass sites ignite with high probability *p_G_* while (b) tree sites ignite with small probability *p*_T_, both from every neighboring burning site. A burning site turns to ash after having the opportunity to spread to neighbouring sites. We repeat this spreading process until no active fires remain. This means fires are ‘instantaneous’ when compared to other timescales.

The first three steps account, in a minimalist way, for vegetation growth, while the final step accounts for consumption of vegetation by fire. The system is typically initialized in a state with no trees. Simulations begin with a 500-time-step burn-in period, to reach the steady state, before we start recording results, which is done only every 50 time steps to avoid autocorrelations.

### Parameter values

#### Timescale

It is helpful define a time step to fix appropriate parameter values. We choose an annual timescale, so that one time step of the model corresponds to one year. To determined the sensitivity of our simulation results to the choice of time step, we also simulate the case in which a time step corresponds to 1/10th of a year and adjust *p_λ_* such that our fastest rate, the rate of grass growth, remains the same. This sampling over different time scales allows us to test the impact, or lack thereof in this case, of having either an instantaneous or probabilistic grass regrowth.

#### Grass growth

Tropical grasses grow back roughly annually, and grass fire intensity saturates after one year (Govender et al., 2006). Therefore, in simulations with an annual time step we use *p_λ_ =* 1, whereas we use *p_λ_ =* 0.1 for a time step equal to a tenth of a year — both corresponding to grass growing back on an annual time scale.

#### Forest propagation

Based on tree maturation times on the order of several decades (Hoffmann et al., 2012a, 2003; Rossatto et al., 2009) and our cell size of 30 × 30 meters, we estimate that a forest propagates at ∼ 1m/year, and therefore set *p_β_ =* 0.03, which is also consistent with empirical observations (Durigan and Ratter, 2006) and previous modeling efforts (Favier et al., 2004).

#### Forest dispersal

Obtaining a quantitative estimate of the rate of generation of new forest patches due to seed dispersal is more difficult. Although long-distance dispersal is common (Fragoso et al., 2003; Nathan, 2006; Nathan et al., 2008), the rate at which new forest patches appear should be much lower than that of propagative forest growth (Clark et al., 1999). Because our model has no memory of how long ago each patch was burned, the probability for creating a new forest patch must incorporate not only the probability of seed dispersal, but also the probability that such a patch avoids fire long enough for a 30 × 30 meter stand of trees to mature and close a canopy. Therefore we set that rate to be at least a factor of 30 lower than propagative forest growth. However, because of the uncertainty in this rate we explore a large range of values for *p_β_* in our simulations.

#### Fire ignition

Typically, any given grassy area in this region burns every 3-7 years (Hoffmann et al., 2012a; Oliveira and Marquis, 2002). We therefore use *p_f_ ∼* 1/(5*L*^2^), which corresponds to a time for a fire to re-occur at a given spatial location of approximately 5 years for a simulation on a lattice of linear size *L* and to a high fire spread probability through grass (*p_G_*).

#### Fire propagation

Because fires spread readily in grasslands but not through tropical forests (Brinck et al., 2017; Cochrane and Laurance, 2002; Hoffmann et al., 2012b; Laurance et al., 2002; Uhl and Kauffman, 1990), we use fire spread probabilities of *p_G_ =* 0.8 - 0.9 in grass and *p_r_ =* 0.1 - 0.2 in trees. However our results are robust as long as *p_G_* and *p_T_* are respectively above and below the site percolation threshold on the square lattice; that is, *p_G_ > p_c_ =* 0.5927…and *p_T_ < p_c_)* (Aharony and Stauffer, 2003). This means that a grass fire can percolate throughout the system, while a tree fire is necessarily finite in spatial extent.

#### Spatial scale

Consistent with our dataset, we assume each grid cell corresponds to an area of 30 × 30m. To check that our results are insensitive to the lattice size, we simulated lattices from 256 × 256 to 768 × 768.

Please see Table 2 for a summary of parameter descriptions and values.

**Table 2:**
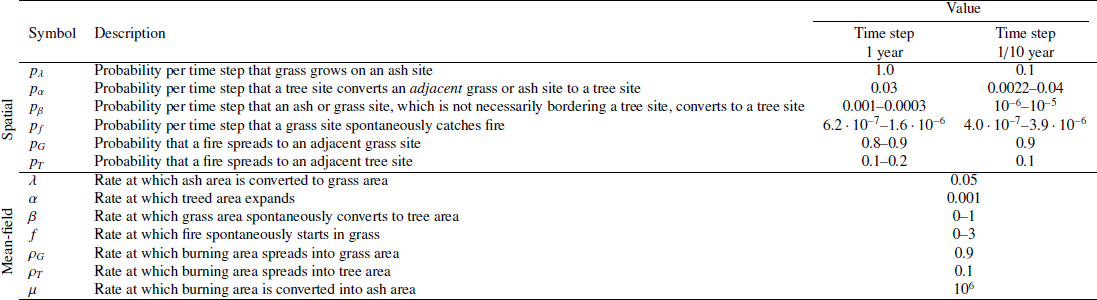
Parameter descriptions and values used for simulations. Note that the numerical between the mean-field and spatial versión of the model should in most cases not be directly compared. For example, values of *p_β_* and *_β_* (and *p_f_* and *f)* do not directly correspond as the impact of the former depends on system size while the latter is defined in an infinite mean-field system.

### Mean-field BGT Model

To develop an analytical understanding of the nature of the savanna-forest transition, we also formulate a mean-field version of our spatial BGT model. In this mean-field version description, the only variables are the global densities of grass, tree, burning and ash sites. The spatial location of sites is are ignored and each site is assumed to neighbor to every other site. The simplicity of the mean-field description allows us to show analytically that the BGT model exhibits both a discontinuous and a continuous transition between a stable fully forested state and a savanna state when basic system parameters, such as fire rate *f* are varied. A discontinuous transition exhibits a sudden macroscopic drop in tree cover with a microscopic change in fire frequency, while the tree cover decreases continuously from a fully-forested state at a continuous transition as a function of model parameters. These transitions mirror the macroscale behavior observed in simulations of the spatial BGT model.

In the mean-field description, spatial degrees of freedom are neglected and the only variables are B, G, T, and A, which now denote the global densities of burning sites, grass, trees, and ash. Following the elemental steps 1-4 of the spatial version outlined above, we show in the appendix that these densities evolve in time as follows:

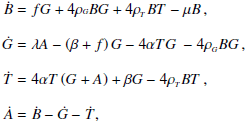

where the overdot denotes the time derivative and conservation of the total area mandates that 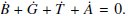
Here, the rates *(λ, α,β, f,ρ_G_,ρ*_r_) correspond to the probabilities that were defined in the 4 steps of the spatial BGT model (see Table 2 for parameter definitions and values). An additional rate—the rate *μ* at which burning sites turn to ash—is needed because the continuous-time dynamics of the mean-field description does not allow for instantaneous events. Instead, the rate at which burning material turns to ash is taken to be arbitrarily large so that it exceeds all other rates in the system (Table 2).

## RESULTS

### Empirical

While the shapes of forest patches can be arbitrarily complicated, there are two clear systematic and quantifiable features of the data. The first is the existence of a scaling relation between the perimeter *P* and area *A* of forest patches. The data for the Cerrado region gives *P ∼ A^γ^* with γ ≈ 0.69 (95% confidence interval [0.68, 0.70]). Similar scaling behavior arises for separate datasets that are partitioned into regions of low (0-1100mm), medium (1100-1800mm), and high annual precipitation (> 1800mm) (Fig. 1E, Table 1) where we find *γ* ≈ 0.70 for the three precipitation cases. Note that for a compact shape, such as a circle or a square, *γ =* 0.5, while for a dendritic shape (in the extreme case, a line) γ ≈ 1. The data indicate that forest patches have a perimeter to area scaling behavior, with *γ* ≈ 0.69, consistently over several orders of magnitude in area. This scaling law also arises in our spatial BGT model (to be discussed in the next section).

### Perimeter to area scaling in the spatial BGT model

Simulations of the spatial BGT model produce a scaling relationship between forest patch perimeters *P* and their areas *A* that is close to what is observed in the Cerrado (Fig. 1E). Using our most realistic parameter values (upper four rows of Table 1), we observe *P ∼ A^γ^,* with an average value of *γ*≈0.70 (95% confidence interval [0.69, 0.71]). Simulations using a wider still range of parameter values and environmental conditions demonstrate the robustness of the scaling relationship: confidence intervals overlap regardless of the annual rainfall in the Cerrado, and regardless of the model parameters in our simulations as long as they lie within a large but realistic range of values (Table 1). In fact, parameter values for fire probability and the relative recruitment rates of trees can be uncertain, and were explored by varying their values by orders of magnitude in our simulations. The consistency between the data and our model prediction for the perimeter-area scaling gives credence to the mechanisms underlying the BGT model. This correspondence between data and theory also allows us to use the BGT model to make non-trivial predictions that are independent of specific model parameter values, such as predicting the fate of individual forest patches, as we show in the next section.

### Fate of forest patches in the spatial BGT model

The results from our model suggest that forest patches converge on an steady-state shape as a result of the balance between expansion by tree propagation and their erosion by fire. We numerically test this hypothesis by simulating the dynamics of individual patches of diverse sizes and shapes in a system that have otherwise reached a steady-state. For this test, we artificially create a forest patch with a given shape and embed it in a much larger steady-state mosaic of grassy and forested areas. The size of this artificial patch ranges from 4 cells up to 10^4^ cells. From initially rectangular patches with aspect ratios in the range of 1:1 to 1:150, we randomly change a fraction of the boundary tree sites to grass sites, in such a way that the overall patch remains connected and also that most of the allowable range of perimeter-area space is sampled (Fig. 2). This allowable range spans the physically realizable portion of the perimeter-area plane where *P ∼ A^γ^*, with 1/2 ≤ *γ* ≤ 1. A few examples of these initial forest patches of small sizes are shown in the periphery of Fig. 2. We simulated at least 10^5^ distinct initial forest patches for a given set of parameter valúes and ran each initial state for five years (5 or 50 time steps (Figs. 2 and C.4 respectively), depending on the timescale). The arrows in Fig. 2 indicate the magnitude and direction of the change in the area and perimeter from the initial state, averaged over many realizations. If a forest patch is fragmented by fire, we track the largest remaining fragment of the original forest, this fragmentation mostly occurs in the top half of the allowable shape range (above the dashed line in Fig. 2). In the bottom half, most forest patches grow in area, but this growth is much slower than the decrease observed in forests that shrink due to fire. Altogether, this experiment allows us to better understand the impact of the shape of a specific forest patch on its stability.

**Figure 2:**
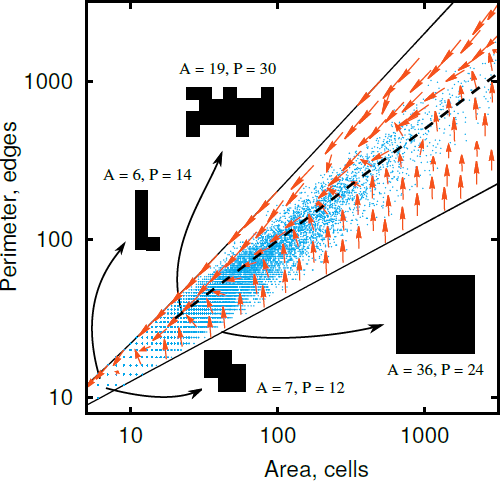
Fate of forest patches. Evolution of forest patches of a given area and perimeter over 5 time steps (red arrows) using the “alternate lattice” set of parameters from Table 1. The 5 time steps therefore provide the expected change in shape over a 5 year period. When a forest is fragmented by fire, we track the largest remaining connected patch. The blue data points correspond to the empirical data. The allowable cone corresponds to all possible forest shapes—dendritic shapes at the upper limit, where the perimeter P is proportional to the area A, and compact shapes at the lower limit, where P ∼ A^1/2^. Some representative starting configurations and their corresponding locations in the perimeter-area plane are shown. The dashed curve P ∼ A^0 7^ corresponds to the locus where the patch evolution is visually the slowest.

### Phase transitions in the spatial BGT model

The state *T =* 1 is an absorbing state of our model; once trees have invaded the system, no more dynamics can occur. In our simulations, we therefore always initiated the system at *T =* 0 to maximize the probability of reaching the savanna steady state and minimize the probability of reaching the absorbing state. For example, for an initial forest cover of *T =* 0.2 and fire probability *p_f_/p_λ_ =* 5 × 10^−6^ (corresponding to the middle of the bistable regime in Fig. 3A), a steady savanna state is reached approximately only 10% of the time while the remaining 90%of simulations reach the absorbing state.

**Figure 3:**
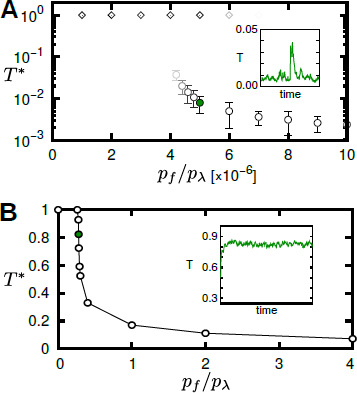
Phase transitions in the spatial BGT model. **(A)** Discontinuous transition for small spontaneous tree growth, *β.* The circles and squares indícate the steady-state values of forest cover, T), as the fire probability, *p_f_*, is varied. Their opacity is related to the probability of reaching this steady state from random initial conditions T(0) = 0.1 (circles) and T(0) = 0.2 (squares). Simulations are on a 512×512 square lattice with *p_β_ =* 2 × 10^−5^ and *p_α_ =* 4 × 10^−3^. *(B)* Continuous transition using no propagative tree spreading *(p_α_ =* 0), *pβ =* 0.02, and *p_λ_ =* 0.1 on a 256×256 square lattice. Standard deviations are smaller than marker size. The insets shows total tree cover as a function of time (T(t)) to illustrate the behaviour of the system at equilibrium. We observe large stochastic fluctuations in tree coverage over time close to the discontinuous transition (panel A) that are not present in the discontinuous case (panel B). The filled green data points in the main figures corresponds to the set of parameters used to produce the time series shown in the insets.

This sensitivity to the initial condition indicates that the system is close to a separatrix in parameter space, where small changes in the initial densities can lead to the system being driven towards different basins of attractions. Correspondingly, there is a discontinuous change in the steady state of the system and a concomitant hysteresis for small changes in the initial condition. These behaviors are consistent with the bistability hypothesized to exist in tropical forests and savannas (Hirota et al., 2011; Staver et al., 2011) and predicted by related models of mixtures of grass and forest (Durrett et al., 2015; Schertzer et al., 2015).

The phenomenology that we predict changes drastically when spontaneous tree growth due to seed dispersion is the dominant contributor to forest growth, a factor that is often omitted in previous models. As illustrated in Fig. 3B, we set an extreme value of *p_α_ =* 0 and examine the behavior for a large value of *p_β_* (10^3^ times larger than for the data in Fig. 3A) that promotes spontaneous tree growth on the grassy areas. In this extreme situation, there is a continuous transition (no discrete jump in tree cover) between the savanna and the fully forested state, as well as much smaller fluctuations in the time series of total amount of observed tree cover (Fig. 3B). This scenario with extremely large *β* leads to an unrealistically rapid growth of trees, but it also illustrates the importance of the *α/β* ratio in shaping the phenomenology of our model. This feature of the BGT model is better studied in its mean-field version.

### Phase transitions in the mean-field BGT model

We can solve for the steady states of the rate equations (1) numerically and gain insights into the nature of the observed phase transitions. These solutions give rise to a cusp bifurcation (Kuznetsov, 2013), leading to three distinct regimes in parameter space: (i) a regime with a single stable solution at a fully forested state *T^*^* = 1; (ii) a regime with a single stable solution which is a mixed-state that corresponds to a savanna (0 < *T^*^ <* 1); (iii) a regime which has two stable solutions, *T*^*^_(1)_ = 1 and 0 < *T*^*^_(2)_ < 1, which are absorbing states for different sets of initial conditions that are demarcated by a separatrix. The transitions between the fully forested and the savanna state are particularly interesting. For sufficiently large values of *β* — i.e., large rate of seed dispersal and ensuing forest patch generation — there is a continuous transition between the forest-dominated state (T = 1) and the savanna state (T < 1) as a function of the fire rate f (Fig. 4A). However, for *β* less than a critical value, *β_c_,* which can only be determined numerically, the tree cover, T, undergoes a discontinuous jump transition instead. This discontinuity is accompanied by both bistability and hysteresis (Fig. 4A and Appendix Fig. E.6). Thus the existence of bistability is determined in part by which tree-growth mechanism—spontaneous growth (β) or neighbor-driven growth *(α*)—is more effective in creating additional trees.

**Figure 4:**
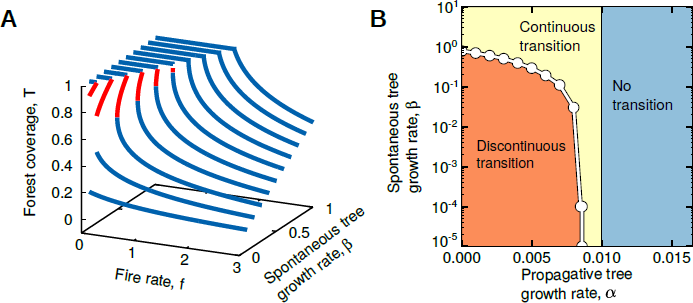
Phase transitions in mean-fleld BGT model. **(A)** Solution of the mean-field model that illustrates the change between a discontinuous and continuous transition. Blue curves are stable steady states, whereas red curves are unstable solutions (missing segments go to the unphysical regime *f <* 0). Note that all the interesting dynamics occur in the realistic regime of slow fire rates f and spontaneous tree growth*β.* **(B)** Nature of the phase transition from the fully treed state (T^*^ = 1) to a mixed state (0 < T^*^ < 1) as a function of propagative and spontaneous tree growth (a and*β* respectively). White markers were obtained by solving for the critical values of *_βc_* that separate the discontinuous and continuous transition regimes. The value of *α_c_* that marks the border of the regime without transition can be derived using the Jacobian of Eqs. (1) (see Appendix). Note that the numerical values of *pβ* and *β* (and *p_f_* and *f*) do not directly correspond as the impact of the former depends on system size while the latter is defined in an infinite mean-field system. The large range of *β* is shown for completeness, but in reality we would not expect*β* to be larger than *α* (uppermost region of figure). In fact, the the rate of propagative growth *α* is by far the most critical mechanism discerning the regime shifts.

## DISCUSSION

Fire spread is an inherently local process, yet is thought to drive vegetation dynamics at a regional scale (Higgins and Scheiter, 2012; Hirota et al., 2011). Localized empirical studies have provided strong support for the hypothesis that different rates of fire spread in savannas and forests (Hoffmann et al., 2012b) largely determine the growth and mortality of tree species (Hoffmann et al., 2009), which in turn can stabilize the savanna-forest boundary (Hoffmann et al., 2012a). By modeling the different fire propagation probabilities in savannas and forests, we are able to reproduce empirically observed patterns at both local scales (the shape and distribution of forest patches) and regional scales (the distribution, or bimodality, of forest coverage). Thus, we connect local mechanisms of fire dynamics, which operate on the scale of meters, to regional patterns of forest and grassland distribution, which operate on the scale of hundreds of kilometers.

Using our spatial model, we showed that commonly observed “edge effects”(Brinck et al., 2017; Cochrane and Laurance, 2002; Laurance et al., 2002; Uhl and Kauffman, 1990) drive the stability and fate of individual forest patches. Importantly, the two competing processes of exposure to fire and propagative growth do not balance because of their very different time scales. For a forest of a given size, having a large perimeter means that it is more likely to be exposed to fire before it receives any benefit from increased tree cover due to propagative growth. Contrarily, if too dense, a forest will not grow sufficiently quickly to offset its exposure to fire, which will lead to an increase in perimeter by encroachment of grassland at the forest edges. The forests that do survive are those that end up with intermediate shapes to ensure enough perimeter growth to offset their exposure to fire.

At the scale of individual forest patches, the basic questions that we addressed were: Will a given forest patch become denser or sparser? Will it grow or will it shrink? Dense shapes (e.g., circular or square forest patches), which correspond to the lower limit of the allowable range in Fig. 2, tend to grow in our simulations, while dendritic shapes that correspond to the upper limit of the range, tend to shrink and/or fragment. A critical feature of this plot is that the locus of points where the observed steady-state perimeter-area relation *P ∼ A^γ^* holds, also corresponds to the locus where the change in the forest perimeter and area is visually the smallest. Thus the most stable forest patches in the spatial BGT model are those that conform to the perimeter-area relation observed in the Brazilian Cerrado (Fig. 1). These results are virtually independent of our choice of parameters. Only extremely unrealistic regimes of *β > α,* i.e. with faster dispersed growth than propagative growth, seem to produce patches with a linear perimeter-area scaling that deviate from the empirical data (see Appendices).

Our spatial model could be a useful tool for resource managers and other stakeholders of tropical forest-savanna systems, because of its potential to predict the fate and risk of collapse of individual forest fragments based on their area and perimeter. This is particularly so due to our model’s lack of sensitivity to the specific parameter values. For example, our model could be used to predict how a specific deformation, such as logging, would affect the stability of a forest patch. However, it is also important to keep in mind that forest edges play a role beyond forest stability. For example, edges are critical to maintaining biodiversity (Turner, 1996). Nonetheless, the validation of our predictions for the fate of specific forest patches, using remotely sensed data with high temporal resolution, will aid in these quantitative predictions.

At the regional scale, we found that our spatial model can exhibit either a sudden discontinuous transition, or a steep but smooth continuous transition, from a fully forested state to a savanna ecotone when parameters change. At this larger scale, the parameterization profoundly affects the results, in contrast, the observations for the fates of individual patches, which were insensitive to specific parameters. Studying the mean-field version of our model allowed us to show that this parameter sensitivity is the result of a cusp bifurcation as a function of three basic parameters—the fire rate *f*, the tree propagation rate *α,* and the spontaneous tree growth rate *β.* The nature of the transition is set by whether tree growth is dominated by propagative forest expansion, which leads to a discontinuous transition, or by tree growth due to seed dispersal, which leads to a continuous transition. This means that factors to increase the recruitment of new forest patches could move the system into a regime with a continuous transition. Such an intervention would be desirable because in such a regime there is less chance of runaway shrinking of forest cover than in the regime with a discontinuous transition. Similarly, animal species that contribute to long-distance seed dispersal (Nathan et al., 2008), especially into grasslands, may play a critical role in limiting discontinuous (or critical) regime shifts.

In previous work, bimodality in the empirical distributions of regional forest cover is often used to hypothesize bistability, and thus discontinuous transitions (criticality), in tropical forests. Our BGT model is consistent with that interpretation since it can also exhibit a discontinuous transition. However, a continuous transition, if sufficiently steep, could give rise to bimodality in forest cover (Fig. 3B). Thus, the empirically observed bimodality is consistent even with our continuous transition, so this bimodality is not necessarily evidence for bistability. This critical distinction between our model and empirical interpretations has implications for how sensitive tropical savanna-forest biomes will be to global change (e.g., Moncrieff et al. (2014)). Empirical studies that quantify the response of various parameters in our model, such as the rate of fire spread, the rate of local tree propagation, etc., to different climatic regimes will allow us to predict the nature of the potential transitions (continuous or discontinuous) between forested and savanna states.

Our parsimonious model comes with several caveats and assumptions. Strictly speaking, we have assumed that mortality when tree sites burn is > 50%, so that even sites at 100% tree cover will fall below the threshold following a fire. In reality, tree mortality due to fire can take on a wide range of values, but our model indirectly accounts for this in two ways. First, a higher value of tree growth due to spreading (*p_α_*) could capture the fast regrowth of peripheral trees that were burned but not completely killed. For this reason, some of our simulations use high *p_α_* values (‘high *α/β* ratio 1’ in Table 1 corresponds to a 4.5 year time scale for tree spreading). Two, one can think of *p_α_* as the product of the probability the trees will burn and the probability they will then die. In this sense, when we varied *p_α_* we effectively varied the fire mortality, and found no change in results. Nonetheless, future models could explicitly consider variability in tree mortality due to factors related to microclimate that influence fire spread and intensity, as well as community composition (Hoffmann et al., 2009) which can vary substantially across the landscape (Pellegrini et al., 2016).

We have also assumed spatial homogeneity in our model. For instance, all grassy areas have an equal probability of burning, which would be the case if there was random ignition across the entire area. In other ecosystems such as coniferous forests, it has been documented that the spread of a single fire can be quite heterogenous due to a number of factors (Turner, 2010). In grasslands, which experience a higher frequency of burning, we expect the heterogeneity in the effect of a single fire ignition and spread to be reduced when multiple fire cycles are considered, because it allows for a greater likelihood that any particular location is burned. On the other hand, local factors, such as underlying topography or even termite mounds (Levick et al., 2010), have the potential to influence fire spread. In these cases, spatial patterns may arise due to the distribution of edaphic features. Similarly, heterogeneity in fire ignition and spread in tropical forests can also be caused by their proximity to roads (Nepstad et al., 2001). Moreover, tree growth (other than edge vs. random) is homogeneous in our model, but in the real world may depend on geographic features, such a topology and proximity to water. Indeed, in the empirical data we observe forest patches whose shape is almost certainly governed by geography (Appendix Fig. A.3). However, while these forests are large in area, they do not contribute many forest patches (i.e., data points) in terms of numbers, so they do not dominate the analysis.

We have assumed binary levels of tree cover (a site is either treed or grass) and homogeneity in their composition. Trees, especially fire adapted ones (Hoffmann et al., 2003; Rossatto et al., 2009), may be sparsely distributed through grassy areas. Our model is consistent with this picture, provided these trees do not significantly impede fire spreading through grass (or if they do it can be incorporated into the fire spread rate *P_G_*). Certainly there is a wide diversity of tree types and demographic characteristics of plants could influence processes such as tree growth and fire spread. However the edge effects that we model are general across species and communities in tropical forest-savanna biomes (Brinck et al., 2017; Cochrane and Laurance, 2002; Laurance et al., 2002; Uhl and Kauffman, 1990). Thus a single tree type (or a homogeneous mix) as modeled here seems adequate to address the questions at hand, but adding more detail (e.g., Berjak and Hearne (2002); Favier et al. (2004); Yassemi et al. (2008)) may be important for other questions.

Finally, testing the generality of our model in different savanna-forest ecotones outside of South America will provide insight into its generality. In African savanna, we may expect fundamentally different dynamics because of the large role that elephants play in determining the distribution of savannas and forests (Asner et al., 2009, 2016). Indeed, both elephants and fire are hypothesized to act in concert to determine alternative stable states of savanna and forest, but unclear whether such edge effects emerge given the ability of elephants to topple large trees (Dublin et al., 1990). In Australia, tree cover can be sensitive to changes in precipitation even in relatively wet conditions.

In conclusion, regardless of model assumptions, we have shown that the low spread rate of fires in forests and its high spread rate in savannas plays a pervasive role in structuring the shape of forest patches across a Neotropical savanna-forest landscape. The ability to infer the stability of an ecosystem from the growth, recruitment, and fire-driven mortality allows us to assess the persistence of both savanna and forest landscapes. Moreover, the ability to infer stability from the shape of forest fragments presents a potential tool for large-scale and rapid assessment of the stability of tropical terrestrial ecosystems. Understanding the key mechanisms at play in shaping forests and their stability will aid in predicting the future of land carbon sinks (Brinck et al., 2017; Pan et al., 2011) and biodiversity (Alroy, 2017; Tilman et al., 1994).

## ACKNOWLEDGMENTS

We thank William F. Laurance, Munik Shrestha and two anonymous reviewers for helpful suggestions. This work has been supported by the Santa Fe Institute, a James S. McDonnell Foundation Postdoctoral Fellowship (LHD), a Santa Fe Institute Omidyar Postdoctoral Fellowship (AMB), a NOAA Climate and Global Change Postdoctoral Fellowship (AFAP) and by the grants DMR-1608211 and 1623243 from the National Science Foundation (UB and SR), the John Templeton Foundation (SR, AMB), and Grant No. 2012145 from the United States Israel Binational Science Foundation (UB).

## REFERENCES

(2016). Category-II: National park, as defined by the International Union for Conservation of Nature (IUCN).

Aharony, A. and Stauffer, D. (2003). Introduction to percolation theory. Taylor & Francis.

Alroy, J. (2017). Effects of habitat disturbance on tropical forest biodiversity. Proceedings of the National Academy of Sciences, 114(23):201611855.

Asner, G. P., Levick, S. R., Kennedy-Bowdoin, T., Knapp, D. E., Emerson, R., Jacobson, J., Colgan, M. S., and Martin, R. E. (2009). Large-scale impacts of herbivores on the structural diversity of african savannas. Proceedings of the National Academy of Sciences, 106(12):4947–4952.

Asner, G. P., Vaughn, N., Smit, I. P., and Levick, S. (2016). Ecosystem-scale effects of megafauna in african savannas. Ecography, 39(2):240–252.

Bak, P., Chen, K., and Tang, C. (1990). A forest-fire model and some thoughts on turbulence. Physics Letters A, 147(5):297–300.

Berjak, S. G. and Hearne, J. W. (2002). An improved cellular automaton model for simulating fire in a spatially heterogeneous savanna system. Ecological modelling, 148(2):133–151.

Brinck, K., Fischer, R., Groeneveld, J., Lehmann, S., De Paula, M. D., Pütz, S., Sexton, J. O., Song, D., and Huth, A. (2017). High resolution analysis of tropical forest fragmentation and its impact on the global carbon cycle. Nature Communications, 8.

Chen, K., Bak, P., and Jensen, M. H. (1990). A deterministic critical forest fire model. Physics Letters A, 149(4):207–210.

Clar, S., Drossel, B., and Schwabl, F. (1996). Forest fires and other examples of self-organized criticality. Journal of Physics: Condensed Matter, 8(37):6803.

Clark, J. S., Silman, M., Kern, R., Macklin, E., and HilleRisLambers, J. (1999). Seed dispersal near and far: patterns across temperate and tropical forests. Ecology, 80(5):1475–1494.

Cochrane, M. A. (2003). Fire science for rainforests. Nature, 421(6926):913–919.

Cochrane, M. A., Alencar, A., Schulze, M. D., Souza, C. M., Nepstad, D. C., Lefebvre, P., and Davidson, E. A. (1999). Positive feedbacks in the fire dynamic of closed canopy tropical forests. Science, 284(5421):1832–1835.

Cochrane, M. A. and Laurance, W. F. (2002). Fire as a large-scale edge effect in amazonian forests. Journal of Tropical Ecology, 18(03):311–325.

Conley, D., Paerl, H., Howarth, R., Boesch, D., Seitzinger, S., Havens, K., Lancelot, C., and Likens, G. (2009). Ecology - controlling eutrophication: Nitrogen and phosphorus. Science, 323(5917):1014–1015.

Dantas, V. d. L., Hirota, M., Oliveira, R. S., and Pausas, J. G. (2016). Disturbance maintains alternative biome states. Ecology Letters, 19(1):12–19.

Drossel, B. and Schwabl, F. (1992). Self-organized critical forest-fire model. Physical Review Letters, 69(11):1629.

Dublin, H. T., Sinclair, A. R., and McGlade, J. (1990). Elephants and fire as causes of multiple stable states in the serengeti-mara woodlands. The Journal of Animal Ecology, pages 1147–1164.

Durigan, G. and Ratter, J. (2006). Successional changes in cerrado and cerrado/forest ecotonal vegetation in western sao paulo state, brazil, 1962-2000. Edinburgh Journal of Botany, 63(1):119–130.

Durrett, R., Zhang, Y., et al. (2015). Coexistence of grass, saplings and trees in the Staver–Levin forest model. The Annals of Applied Probability, 25(6):3434–3464.

Favier, C., Chave, J., Fabing, A., Schwartz, D., and Dubois, M. A. (2004). Modelling forest-savanna mosaic dynamics in man-influenced environments: effects of fire, climate and soil heterogeneity. Ecological Modelling, 171(1):85–102.

Fragoso, J., Silvius, K. M., and Correa, J. A. (2003). Long-distance seed dispersal by tapirs increases seed survival and aggregates tropical trees. Ecology, 84(8):1998–2006.

Govender, N., Trollope, W. S., and Van Wilgen, B. W. (2006). The effect of fire season, fire frequency, rainfall and management on fire intensity in savanna vegetation in south africa. Journal of Applied Ecology, 43(4):748–758.

Grassberger, P. (2002). Critical behaviour of the Drossel-Schwabl forest fire model. New Journal of Physics, 4(1):17.

Hansen, M. C., Potapov, P. V., Moore, R., Hancher, M., Turubanova, S., Tyukavina, A., Thau, D., Stehman, S., Goetz, S., Loveland, T., et al. (2013). High-resolution global maps of 21st-century forest cover change. Science, 342(6160):850–853.

Higgins, S. I. and Scheiter, S. (2012). Atmospheric co2 forces abrupt vegetation shifts locally, but not globally. Nature, 488(7410):209–212.

Hirota, M., Holmgren, M., Van Nes, E. H., and Scheffer, M. (2011). Global resilience of tropical forest and savanna to critical transitions. Science, 334(6053):232–235.

Hoffmann, W. A., Adasme, R., Haridasan, M., T de Carvalho, M., Geiger, E. L., Pereira, M. A., Gotsch, S. G., and Franco, A. C. (2009). Tree topkill, not mortality, governs the dynamics of savanna–forest boundaries under frequent fire in central brazil. Ecology, 90(5):1326–1337.

Hoffmann, W. A., Geiger, E. L., Gotsch, S. G., Rossatto, D. R., Silva, L. C., Lau, O. L., Haridasan, M., and Franco, A. C. (2012a). Ecological thresholds at the savanna-forest boundary: how plant traits, resources and fire govern the distribution of tropical biomes. Ecology letters, 15(7):759–768.

Hoffmann, W. A., Jaconis, S. Y., Mckinley, K. L., Geiger, E. L., Gotsch, S. G., and Franco, A. C. (2012b). Fuels or microclimate? understanding the drivers of fire feedbacks at savanna–forest boundaries. Austral Ecology, 37(6):634–643.

Hoffmann, W. A., Orthen, B., and Nascimento, P. K. V. d. (2003). Comparative fire ecology of tropical savanna and forest trees. Functional Ecology, 17(6):720–726.

Kuznetsov, Y. A. (2013). Elements of applied bifurcation theory, volume 112. Springer Science & Business Media.

Laurance, W. F., Lovejoy, T. E., Vasconcelos, H. L., Bruna, E. M., Didham, R. K., Stouffer, P. C., Gascon, C., Bierregaard, R. O., Laurance, S. G., and Sampaio, E. (2002). Ecosystem decay of amazonian forest fragments: a 22-year investigation. Conservation Biology, 16(3):605–618.

Laurance, W. F. and Yensen, E. (1991). Predicting the impacts of edge effects in fragmented habitats. Biological Conservation, 55(1):77–92.

Levick, S. R., Asner, G. P., Chadwick, O. A., Khomo, L. M., Rogers, K. H., Hartshorn, A. S., Kennedy-Bowdoin, T., and Knapp, D. E. (2010). Regional insight into savanna hydrogeomorphology from termite mounds. Nature communications, 1:65.

Moncrieff, G. R., Scheiter, S., Bond, W. J., and Higgins, S. I. (2014). Increasing atmospheric co2 overrides the historical legacy of multiple stable biome states in africa. New Phytologist, 201(3):908–915.

Mumby, P. J., Hastings, A., and Edwards, H. J. (2007). Thresholds and the resilience of caribbean coral reefs. Nature, 450(7166):98–101.

Nathan, R. (2006). Long-distance dispersal of plants. Science, 313(5788):786–788.

Nathan, R., Schurr, F. M., Spiegel, O., Steinitz, O., Trakhtenbrot, A., and Tsoar, A. (2008). Mechanisms of long-distance seed dispersal. Trends in ecology & evolution, 23(11):638–647.

Nepstad, D., Carvalho, G., Barros, A. C., Alencar, A., Capobianco, J. P., Bishop, J., Moutinho, P., Lefebvre, P., Silva, U. L., and Prins, E. (2001). Road paving, fire regime feedbacks, and the future of amazon forests. Forest ecology and management, 154(3):395–407.

Nepstad, D. C., Verssimo, A., Alencar, A., Nobre, C., Lima, E., Lefebvre, P., Schlesinger, P., Potter, C., Moutinho, P., Mendoza, E., et al. (1999). Large-scale impoverishment of amazonian forests by logging and fire. Nature, 398(6727):505–508.

Oliveira, P. S. and Marquis, R. J. (2002). The Cerrados of Brazil: Ecology and Natural History of a Neotropical Savanna. Columbia University Press.

Pan, Y., Birdsey, R. A., Fang, J., Houghton, R., Kauppi, P. E., Kurz, W. A., Phillips, O. L., Shvidenko, A., Lewis, S. L., Canadell, J. G., et al. (2011). A large and persistent carbon sink in the world’s forests. Science, 333(6045):988–993.

Pellegrini, A. F., Franco, A. C., and Hoffmann, W. A. (2016). Shifts in functional traits elevate risk of fire-driven tree dieback in tropical savanna and forest biomes. Global change biology, 22(3):1235–1243.

Pueyo, S., De Alencastro Graça, P. M. L., Barbosa, R. I., Cots, R., Cardona, E., and Fearnside, P. M. (2010). Testing for criticality in ecosystem dynamics: the case of amazonian rainforest and savanna fire. Ecology Letters, 13(7):793–802.

Romme, W. H. and Despain, D. G. (1989). Historical perspective on the yellowstone fires of 1988. BioScience, 39(10):695–699.

Rossatto, D. R., Hoffmann, W. A., and Franco, A. C. (2009). Differences in growth patterns between co-occurring forest and savanna trees affect the forest-savanna boundary. Functional Ecology, 23(4):689–698.

Scheffer, M., Carpenter, S., Foley, J. A., Folke, C., and Walker, B. (2001). Catastrophic shifts in ecosystems. Nature, 413(6856):591–596.

Schertzer, E., Staver, A., and Levin, S. (2015). Implications of the spatial dynamics of fire spread for the bistability of savanna and forest. Journal of Mathematical Biology, 70(1-2):329–341.

Smith, V. H. and Schindler, D. W. (2009). Eutrophication science: where do we go from here? Trends in Ecology & Evolution, 24(4):201–207.

Staver, A. C., Archibald, S., and Levin, S. A. (2011). The global extent and determinants of savanna and forest as alternative biome states. Science, 334(6053):230–232.

Tilman, D., May, R. M., Lehman, C. L., and Nowak, M. A. (1994). Habitat destruction and the extinction debt. Nature, 371:65–66.

Turner, I. (1996). Species loss in fragments of tropical rain forest: a review of the evidence. Journal of applied Ecology, pages 20–209.

Turner, M. G. (2005). Landscape ecology: what is the state of the science? Annu. Rev. Ecol. Evol. Syst., 36:319–344.

Turner, M. G. (2010). Disturbance and landscape dynamics in a changing world. Ecology, 91(10):2833–2849.

Uhl, C. and Kauffman, J. B. (1990). Deforestation, fire susceptibility, and potential tree responses to fire in the eastern amazon. Ecology, 71(2):437–449.

Yassemi, S., Dragićević, S., and Schmidt, M. (2008). Design and implementation of an integrated gis-based cellular automata model to characterize forest fire behaviour. ecological modelling, 210(1):71–84.

